# Accurate quantification of spliced and unspliced transcripts for single-cell RNA sequencing with tidesurf

**DOI:** 10.1101/2025.01.28.635274

**Authors:** Jan T. Schleicher, Manfred Claassen

## Abstract

**Motivation:** Single-cell RNA sequencing (scRNA-seq) allows for the detailed analysis of dynamic cellular processes. In particular, this has been enabled by the estimation of RNA velocity, the derivative of gene expression, from separate count matrices for different splice states, which provides information about a cell’s immediate future even in snapshot data. Useful velocity estimates strongly depend on accurate counts for spliced and unspliced transcripts. Velocyto remains the standard tool for spliced and unspliced mRNA molecule quantification. However, despite considerable advances in scRNA-seq protocols, velocyto has not been updated to account for peculiarities of new protocols, such as popular approaches based on 5’ chemistry.

**Results:** To address this shortcoming, we present tidesurf, a command line tool for the quantification of spliced and unspliced transcript molecules from scRNA-seq libraries. Employing it on four different publicly available 10x Genomics Chromium datasets, we show the accuracy on various datasets generated with either 3’ or 5’ chemistry, whereas velocyto’s results are highly erroneous for the latter. Considering broader applicability, our results highlight tidesurf as a potential replacement for velocyto.

**Availability and implementation:** A Python implementation of tidesurf is available from PyPI and at github.com/janschleicher/tidesurf. Code for reproducing the analyses is available at github.com/janschleicher/tidesurf projects.

## 1 Introduction

Over the last decade, single-cell RNA sequencing (scRNA-seq) has established itself as a method routinely used to study dynamic processes of cells in development, differentiation, diseases, and immune responses [Griffiths et al., 2018, Nayak and Hasija, 2021].

The first droplet-based scRNA-seq protocols were based on capturing the poly-A tail at the 3’ end of transcripts through oligo-dT nucleotide-coated gel beads. More recently, additional protocols have been developed to capture the 5’ end of mRNA molecules, enabling, for example, the concurrent analysis of T or B cell receptor repertoires. In both cases, in the following denoted 5’ and 3’ scRNA-seq, sequencing of the gene expression library results in two separate read files, with the first read containing the cell barcode (CBC) and unique molecular identifier (UMI) and the second read containing the captured mRNA sequence. The resulting data is used to generate a cell-by-gene read count matrix. This requires read alignment to a reference genome and transcript quantification. The scRNA-seq protocols are implemented on the 10x Genomics Chromium platform, which further provides the Cell Ranger software based on the STAR aligner to achieve the alignment and quantification tasks [Zheng et al., 2017, Dobin et al., 2012]. While Cell Ranger considers both exonic and intronic read alignments, it does not quantify them separately but instead produces a single gene expression matrix.

La Manno et al. [2018] first described how a separate quantification of those reads can be leveraged for the analysis of dynamic cellular processes from snapshot data. Their tool, velocyto, produces distinct count matrices for spliced, unspliced, and ambiguously assigned mRNA molecules, based on the Cell Ranger Binary Alignment Map (BAM) files. Other tools such as alevin-fry [He et al., 2022] and kallisto-bustools [Melsted et al., 2021], based on the pseudo-aligners salmon [Patro et al., 2017] and kallisto [Patro et al., 2017], respectively, are also capable of producing separate count matrices, using the raw sequencing FASTQ files as input. From these matrices, so-called RNA velocity, the temporal derivative of gene expression, can be estimated through a simple differential equation model.

While the estimation of RNA velocity has been refined by many other methods such as scVelo [Bergen et al., 2020], it is usually still based on spliced and unspliced counts computed with velocyto. This is potentially problematic as velocyto has not been updated since the beginning of 2019, despite the significant progress in scRNA-seq protocols. For example, it does not offer a mode for the analysis of 5’ scRNA-seq data, resulting in severe quantification errors when used on such datasets [Klingler et al., 2025].

Here, we present a Tool for IDentification and Enumeration of Spliced and Unspliced Read Fragments (tidesurf) from the Cell Ranger BAM files which, unlike velocyto, was developed for both 3’ and 5’ data generated with 10x Genomics Chromium. We used tidesurf to reanalyze four publicly available scRNA-seq datasets (two for each protocol) to demonstrate its utility. Comparing the results to the analysis of the same datasets with velocyto and alevin-fry, we demonstrated its accuracy concerning the quantification of spliced, unspliced, and ambiguous counts. Furthermore, we could show that RNA velocities estimated from spliced and unspliced counts computed with tidesurf were in line with previous results.

### 2 Methods

### 2.1 Implementation

We developed tidesurf, a command line tool for quantifying spliced and unspliced transcripts, implemented in Python. The tool takes a Gene Transfer Format (GTF) file with gene and transcript annotations for the reference genome used for read alignment and a Cell Ranger [Zheng et al., 2017] output directory as input (Fig. 1A).

**Fig. 1.**
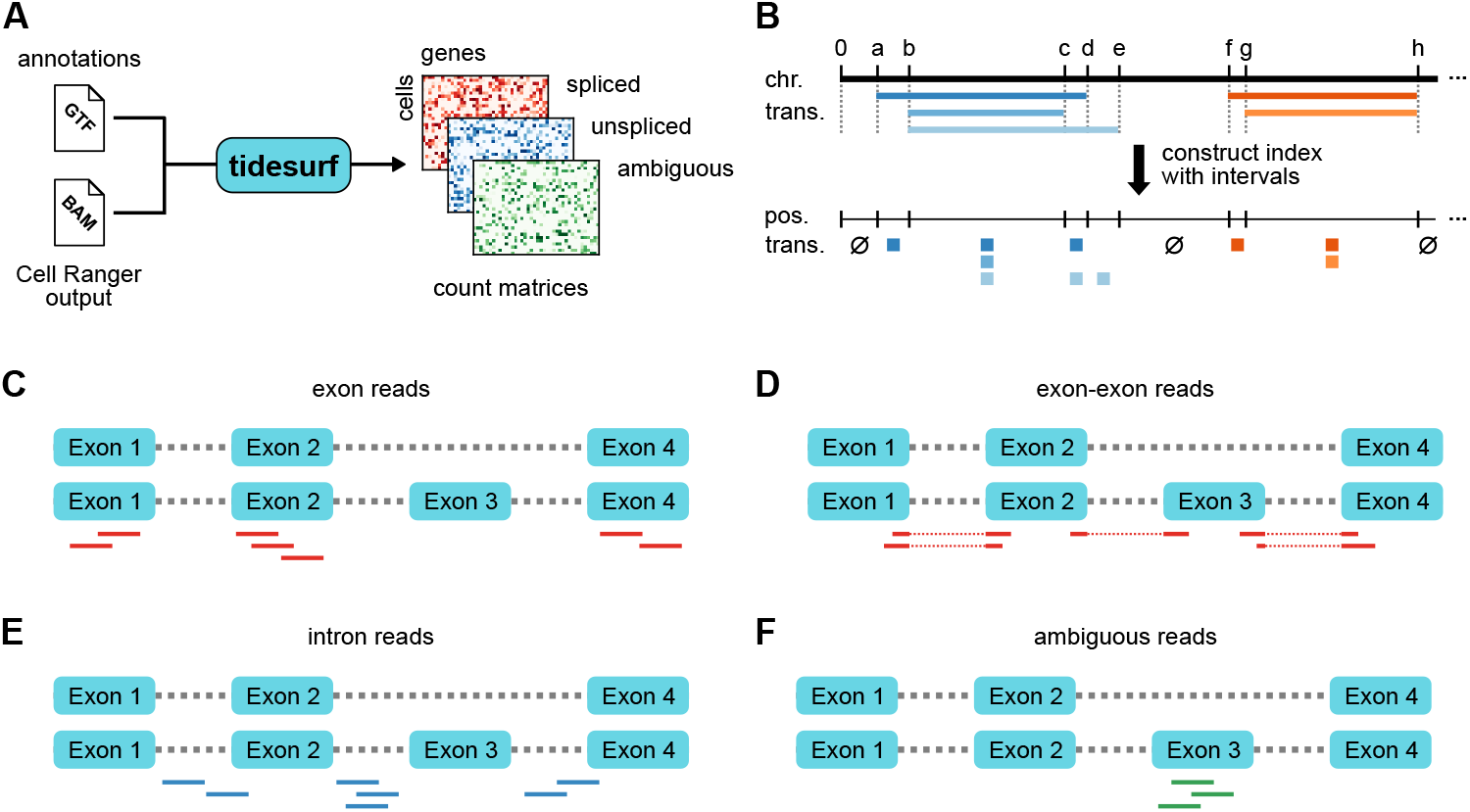
Overview of the tidesurf algorithm. **A** Tidesurf takes a GTF file with transcript annotations and the Cell Ranger output, containing the alignments in BAM format, as input and produces count matrices for spliced, unspliced, and ambiguous mRNA molecules. **B** The annotated genome is used to construct an index. To this end, a list of intervals represented by their genomic starting position is constructed, where for each interval, the set of overlapping transcripts is stored. **C-F** Examples of reads for the different read types used in tidesurf. Two example transcripts with different final exon compositions are shown. Exon reads overlap with exons for all transcripts for a gene (**C**), while exon-exon reads span junctions between exons (**D**). Intron reads are those aligned to introns for all transcripts (**E**), while ambiguous reads map to exons for some of the transcripts, and to introns for others (**F**).

In the first step, a transcript index is built from the GTF file to efficiently retrieve transcripts overlapping with aligned sequencing reads. To this end, all transcripts on a particular strand (plus or minus) and chromosome in the GTF file and their exons are inserted into a sorted list of intervals (Fig. 1B). Each entry in the list consists of the start position of the respective interval and the set of transcripts contained therein. Hence, each interval start corresponds to the start of one or multiple transcripts or the base directly after one or multiple transcripts.

In the second step, the Binary Alignment Map (BAM) file produced by Cell Ranger is processed. The aligned reads are processed iteratively. Unmapped reads, those that have a mapping quality below 255 (i.e. not confidently mapped to a single locus), or reads that do not have a cell barcode (CBC) or unique molecular identifier (UMI) tag are discarded. All other reads are handled individually to determine the gene of origin and splicing status.

First, all transcript annotations overlapping the read are extracted from the transcript index using a binary search for the genomic start and end positions of the read alignment. Since the list of intervals and transcripts contained therein is sorted, this allows us to obtain all relevant transcripts. Here, we only take the strand into account that has the correct orientation relative to the read alignment. For 10x Genomics Chromium, this is the sense orientation for 3’ and the antisense orientation for 5’ protocols. Next, we determine for each overlapping transcript if the read maps to an exonic or intronic region (Fig. 1C-F). This is done by iterating over the exons and introns of the transcript and summing up the respective base overlaps, taking the Concise Idiosyncratic Gapped Alignment Report (CIGAR) string of the aligned read into account. Furthermore, we keep track of the number of exons aligned with the read. For each transcript, we determine the read type based on the following criteria. If the number of bases not overlapping with exons is smaller than a threshold (default: 5), the read is classified as exonic (Fig. 1C) or as spanning an exon-exon junction (Fig. 1D), depending on whether the number of exons is 1 or greater than 1. On the other hand, if the number of bases overlapping with introns is greater than or equal to that threshold, the read is classified as intronic with respect to the transcript (Fig. 1E). Finally, we determine the overall type of a read based on all transcripts. If all transcripts belong to the same gene, a read is classified as spanning an exon-exon junction if that annotation is present. Otherwise, it is classified as either intronic or exonic if the same type was inferred for all transcripts or ambiguous if the type differs between transcripts (Fig. 1F). The rationale behind this is that a read aligning to a region that is an exon for one splice isoform, but an intron for another one cannot be unambiguously assigned unless it spans an exon-exon junction for one of the transcripts. This behavior only occurs when an intron is spliced out.

If the transcripts belong to multiple genes, the default is to discard the read. Alternatively, a read type can be assigned per gene as described above. In that case, it is weighted by the inverse of the number of genes for downstream processing.

In the third step, CBCs and UMIs are deduplicated. For each CBC/UMI/gene/read type combination, we compute its read support. After filtering out read types with very low counts and percentages, we retain only one read type for each barcode/UMI/gene combination. Here, we keep the first read type, where the sort order is intronic *<* exon-exon junction *<* ambiguous *<* exonic. Then, we map the read type to a splice type, where intronic maps to unspliced, ambiguous stays ambiguous, and the two exonic read types map to spliced.

Then, multi-mapped UMIs are resolved. Following Cell Ranger’s gene expression algorithm [10x Genomics, 2024], we assign a CBC/UMI combination to the gene with the highest read support. Ties for maximal read support are discarded.

Finally, spliced, unspliced, and ambiguous UMIs are counted for each cell and gene, resulting in three separate count matrices which are combined into and saved as an AnnData object [Virshup et al., 2024].

### 2.2 Data retrieval

For the mouse pancreatic endocrinogenesis dataset at embryonic day 15.5 [Bastidas-Ponce et al., 2019], FASTQ files were obtained from the Gene Expression Omnibus (GEO, sample accession number GSM3852755). Processed data was obtained from the scVelo (v0.3.3) pancreas dataset.

For the dataset of developing mouse retinal neurons [Lo Giudice et al., 2019], FASTQ files for batch F2 were downloaded from GEO (sample accession number GSM3466902). Processed data was downloaded from the dataset link provided by the authors of UniTVelo [Gao et al., 2022] on the UniTVelo GitHub repository.

Raw FASTQ files and processed data of human peripheral B cells [Stewart et al., 2021] were obtained from ArrayExpress (accession number E-MTAB-9544).

BAM files and processed data of human alloreactive T cells [Fu et al., 2023] were accessed from GEO (series accession number GSE252994). The BAM files were converted to FASTQ using 10x Genomics `bamtofastq`.

### 2.3 Cell Ranger

Raw reads were aligned to the respective annotated reference genome mouse mm10 (GENCODE vM23/Ensembl98) or human GRCh38 (GENCODE v32/Ensembl98), obtained from the Cell Ranger 2020-A reference packages, using 10x Genomics Cell Ranger (v7.1.0) count with automatic chemistry detection [Zheng et al., 2017].

### 2.4 Alevin-fry

Spliced + intronic (splici) references were generated from the Cell Ranger reference transcriptomes with pyroe (v0.9.3) according to the read length of the respective study (Bastidas-Ponce et al. [2019]: 151 bases, Lo Giudice et al. [2019]: 100 bases, Stewart et al. [2021]: 100 bases, and Fu et al. [2023]: 91 bases (samples MJ001, MJ002, MJ003, MJ016, and MJ017) or 101 bases (samples MJ005, MJ006, MJ007, MJ008, MJ009,

MJ018, and MJ019)) with a flank trim length of 5 bases. Transcriptome indices were built with salmon (v1.10.2) `index` [Patro et al., 2017]. Reads were mapped against the respective index with `salmon alevin` with library type ISR for 3’ data and ISF for 5’ data. For quantification of spliced and unspliced reads with alevin-fry (v0.8.2), permit lists were generated using `generate-permit-list` with expected orientation `fw` for 3’ data and `rc` for 5’ data, and barcodes were corrected with `collate` [He et al., 2022]. Finally, `quant` was used for quantification. The results were transformed into an AnnData object using `load_fry` from pyroe.

### 2.5 Velocyto

To obtain spliced and unspliced counts with velocyto (v0.17.17), the Cell Ranger output directories were processed using `run10x` [La Manno et al., 2018] with the respective annotation GTF file.

### 2.6 Tidesurf

For quantification of spliced and unspliced transcripts with tidesurf (v0.1.0), the Cell Ranger output directories were processed using default parameters and the respective gene annotation GTF files from the Cell Ranger references. For 3’ data, the orientation was specified as `sense`, while it was set to `antisense` for 5’ datasets.

### 2.7 Data processing and RNA velocity analysis

All datasets were processed using scanpy (v1.10.4) [Wolf et al., 2018] and scVelo (v0.3.3) [Bergen et al., 2020]. For comparisons of counts per cell (total or splice state-specific), the processed reference datasets and the count matrices obtained from Cell Ranger, velocyto, alevin-fry, and tidesurf were filtered to contain only common cells (Table 1).

**Table 1.** Cell numbers in the analyzed datasets.

| Dataset | Publication | Number of cells |  |
| --- | --- | --- | --- |
|  |  | Reference | Common |
| Pancreas | Bastidas-Ponce et al. [2019] | 3,696 | 3,696 |
| Retina | Lo Giudice et al. [2019] | 2,726 | 2,635 |
| B cells | Stewart et al. [2021] | 8,771 | 7,312 |
| T cells | Fu et al. [2023] | 51,504 | 50,448 |

For downstream RNA velocity analysis, the datasets were additionally subset to common genes. Genes with less than 20 counts (spliced and unspliced) were filtered out. The gene expression matrix as well as the spliced and unspliced counts were normalized so that the total counts per cell were equal to the median total counts before normalization. Highly variable genes were determined with `scvelo.pp.filter_genes_dispersion`, using the `seurat` flavor and the top 2,000 genes. The expression matrix was log(*x* + 1)-transformed. The first 30 principal components from the respective reference dataset were used to compute a neighborhood graph with 30 neighbors. Based on this neighborhood graph, moments of spliced and unspliced counts were calculated and velocities were computed using scVelo’s stochastic model. Finally, velocities were projected onto the 2-dimensional UMAP embedding of the respective reference dataset for visualization.

## 3. Results

To demonstrate the utility of tidesurf, we analyzed four publicly available datasets generated with the 10x Genomics Chromium technology.

The pancreatic endocrinogenesis [Bastidas-Ponce et al., 2019] and the developing retina datasets [Lo Giudice et al., 2019] were generated with the 3’ chemistry, while the peripheral B cell [Stewart et al., 2021] and alloreactive T cell datasets [Fu et al., 2023] were generated with the 5’ chemistry. For all four datasets, we compared the read count output of tidesurf to the Cell Ranger expression matrix and quantifications obtained with the tools velocyto and alevin-fry. Further, we compared the RNA velocity estimates based on the quantifications by the competing approaches.

### 3.1 Counts obtained with tidesurf recapitulate the Cell Ranger quantification

First, we assessed the total read count outputs. For the 3’ datasets, the distribution of total counts per cell (sum of spliced, unspliced, and ambiguous counts) was very similar between all three methods and Cell Ranger, with velocyto showing slightly lower total counts (Fig. 2A). In contrast, velocyto total counts were considerably lower than those of alevin-fry, tidesurf, and Cell Ranger for the 5’ datasets, especially for the Stewart et al. [2021] data. The same trend was observed for the difference of total counts per cell of velocyto, alevin-fry, and tidesurf from Cell Ranger (Fig. 2B). The median difference of tidesurf total counts from Cell Ranger was less than 100 for all datasets except for the T cell data, where it was slightly higher at around 280. For alevin-fry, the absolute median difference ranged between approximately 200 on the 3’ datasets and almost 800 on the B cell data. Velocyto’s median difference was around 500 for the retina data, around 1,000 for the pancreas and T cell datasets, and above 3,000 for the B cell data. The counts showed a high Pearson correlation with the Cell Ranger counts of the same gene for alevin-fry and tidesurf on all four datasets (median correlation always above 0.94), while for velocyto, this was only the case for the 3’ datasets (Fig. 2C). Similarly, cell-wise cosine similarities of the counts with the Cell Ranger expression matrix were high on all four datasets for alevin-fry and tidesurf (median similarity above 0.95; Fig. 2C). Even so, cosine similarities to Cell Ranger tended to be slightly higher for tidesurf than for alevin-fry, particularly on the 5’ datasets. In contrast, velocyto had a much lower similarity on the 5’ datasets, with a median similarity of around 0.8 on the T cell and below 0.1 on the B cell data. The notable discrepancies for velocyto on 5’ scRNA-seq data are not surprising as velocyto was developed before the establishment of 10x Genomics’ 5’ protocols and has not been updated since. This was analyzed and described in detail by Klingler et al. [2025].

**Fig. 2.**
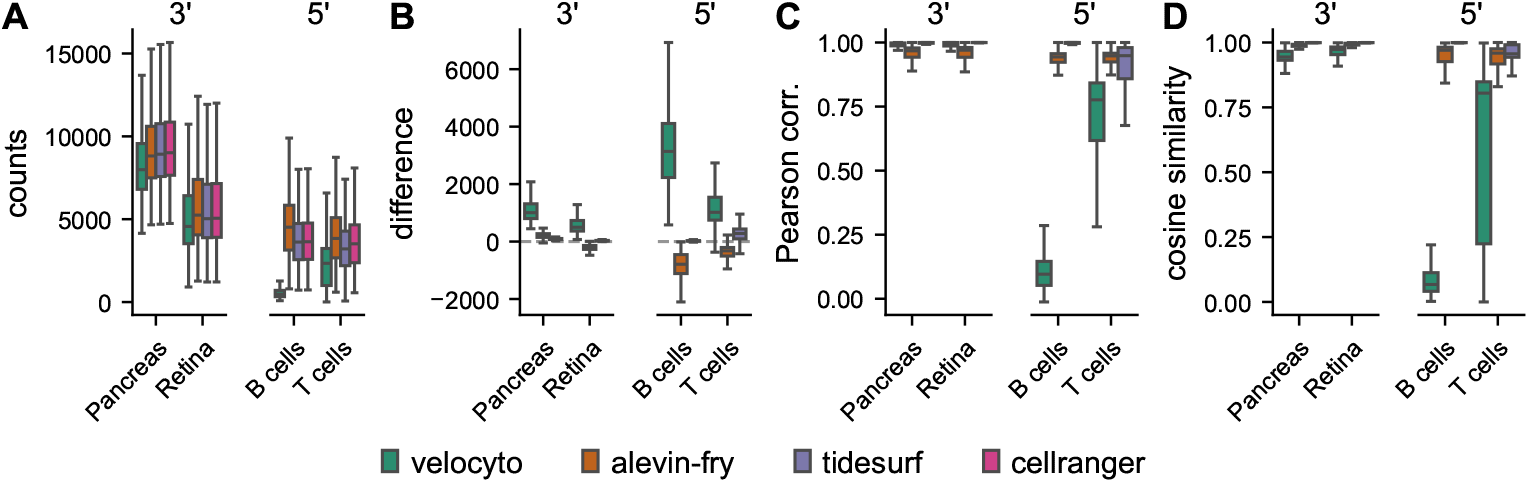
Comparison of counts across methods. **A** Total counts per cell (sum of spliced, unspliced, and ambiguous for velocyto, alevin-fry, and tidesurf) per dataset. **B** Difference of total counts per cell from Cell Ranger counts. **C** Per-gene Pearson correlation of counts with Cell Ranger counts. **D** Per-cell cosine similarity of counts with Cell Ranger counts.

### 3.2 Spliced, unspliced, and ambiguous counts are similar between tidesurf and alevin-fry

Furthermore, we separately compared the total spliced, unspliced, and ambiguous counts per cell before normalization. For the 3’ datasets, the distributions of total spliced counts were similar between all three methods (Fig. 3A). In contrast, velocyto produced considerably lower spliced counts than alevin-fry and tidesurf. The distributions of unspliced counts were comparable between all tools on all four datasets, regardless of the assay chemistry (Fig. 3B). Alevin-fry tended to produce higher ambiguous counts than velocyto and tidesurf (Fig. 3C). Accordingly, the total counts differences per cell from the tidesurf quantification were low for spliced counts for alevin-fry on all four datasets, but more pronounced for velocyto on the 5’ datasets (Fig. S1A). On the pancreas and retina datasets, velocyto quantified more molecules as unspliced than tidesurf (Fig. S1B) and achieved similar total ambiguous counts (Fig. S1C), whereas alevin-fry produced lower total spliced counts and higher ambiguous counts on all four datasets. This difference may stem from the different resolution strategies for UMI splice types. In contrast to velocyto and tidesurf, alevin-fry bases its assignment of a UMI to a splice type on a majority vote of its corresponding reads.

**Fig. 3.**
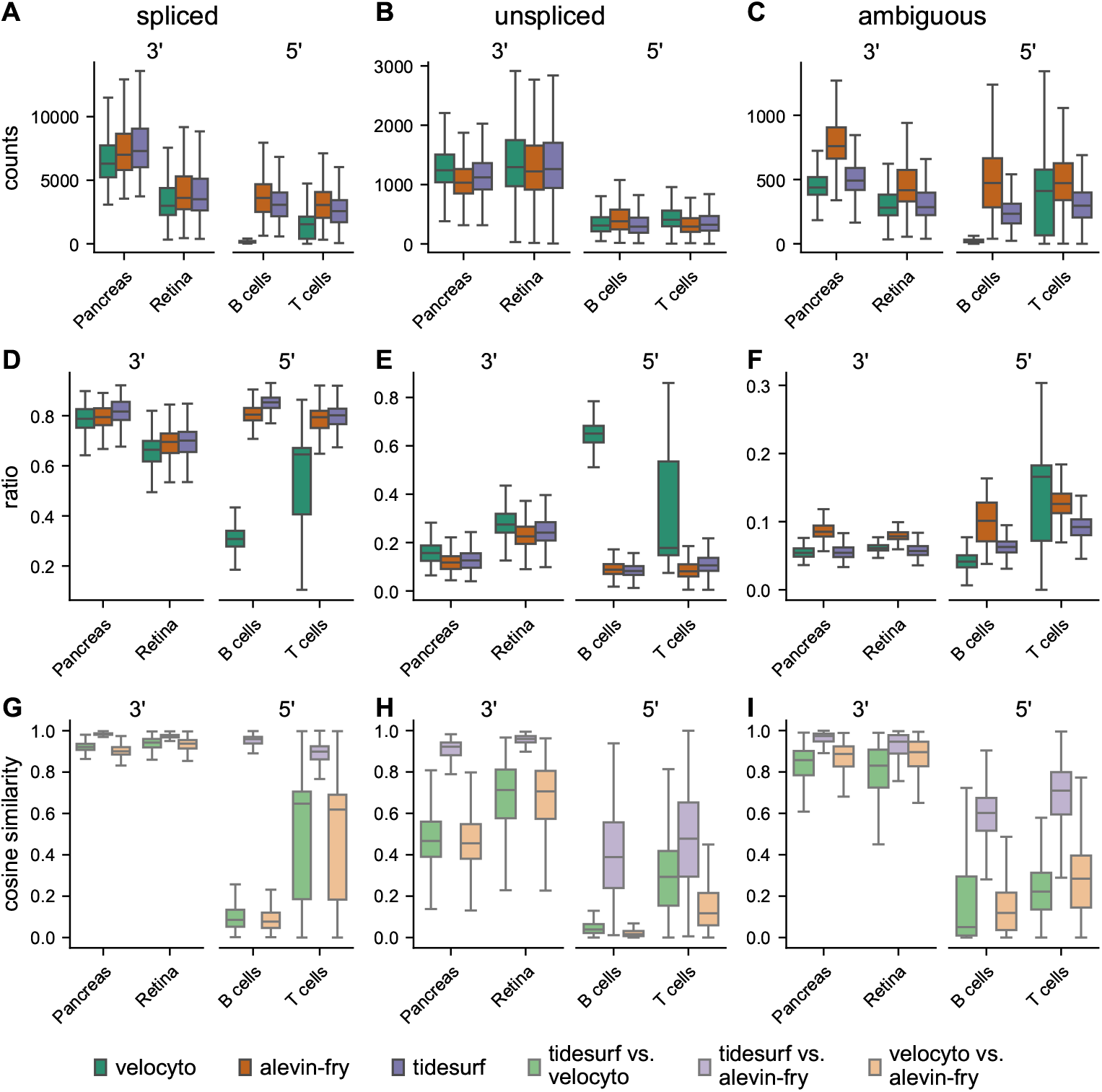
Comparison of spliced, unspliced, and ambiguous counts across methods. **A-C** Total spliced (**A**), unspliced (**B**), and ambiguous (**C**) counts per cell. **D-F** Ratio of spliced (**D**), unspliced (**E**), and ambiguous (**F**) counts out of total counts per cell. **G-I** Cosine similarity of spliced (**G**), unspliced (**H**), and ambiguous (**I**) counts per cell between methods.

Similarly, the ratios of spliced (Fig. 3D), unspliced (Fig. 3E), and ambiguous counts (Fig. 3F) per cell out of all counts were in a comparable range for all methods on the 3’ datasets, while velocyto had much lower proportions of spliced and much higher proportions of unspliced molecules for the 5’ datasets.

For a direct, pairwise comparison between methods, we investigated the per-cell cosine similarities and per-gene Pearson correlations of spliced, unspliced, and ambiguous counts, respectively. For this analysis, we only considered genes present for all three methods. On the 3’ datasets, the median cosine similarity of spliced counts between tidesurf and alevin-fry was close to 1, and above 0.9 for the comparisons between tidesurf and velocyto, and between velocyto and alevin-fry (Fig. 3G). In contrast, the median similarity between tidesurf and velocyto and between velocyto and alevin-fry was below 0.1 for the B cell data and between 0.6 and 0.65 for the T cell data. The median Pearson correlation was above 0.95 for all pairwise comparisons on the pancreas and retina datasets, and around 0.9 between tidesurf and alevin-fry on the B cell and T cell datasets (Fig. S1D). Like the cosine similarity, the Pearson correlation between velocyto’s spliced counts and those of the other two methods was also considerably lower on the 5’ datasets, especially on the B cell data. For unspliced counts, the median cosine similarity between tidesurf and alevin-fry was above 0.9 on the 3’ datasets, and around 0.4 and 0.5 on the B cell and T cell data, respectively, while velocyto had a lower similarity to both tidesurf and alevin-fry for all datasets, with particularly low values for the data from Stewart et al. [2021] (median below 0.05, Fig. 3H). A similar trend was observed for gene-wise Pearson correlations on the 5’ datasets, while velocyto had slightly higher correlations to tidesurf than the other two comparisons on the 3’ data (Fig. S1E). Ambiguous counts had a median cosine similarity above 0.83 and a median correlation between 0.6 and 0.8 on the pancreas and retina datasets for all comparisons, a median cosine similarity between tidesurf and alevin-fry of around 0.6 and 0.7, and a median correlation between tidesurf and alevin-fry of 0.6 and 0.5, on the B cell and T cell data, respectively (Fig. 3I, Fig. S1F). The similarity and correlation between velocyto and the other two methods were considerably lower. As described above, the considerable differences between velocyto and the other two tools on the 5’ datasets were expected since velocyto’s quantification is highly erroneous for these protocols [Klingler et al., 2025].

There are multiple plausible reasons for the higher differences between alevin-fry and tidesurf on the 5’ datasets compared to the 3’ datasets. First, the 3’ datasets stem from murine samples while the 5’ datasets originate from human donors. The human genome contains more repetitive sequences than the murine genome [Lu et al., 2020], which could lead to differences in the (pseudo-)alignments of Cell Ranger and alevin-fry, especially for intronic sequences. Furthermore, the total unspliced counts were considerably lower on the two 5’ datasets, rendering small absolute differences relatively more important. Finally, alevin-fry’s quantification of total counts differed slightly more from the Cell Ranger output than tidesurf’s (Fig. 2B-D).

### 3.3 RNA velocities and their interpretation are similar between quantification methods

Finally, we computed RNA velocities based on the spliced and unspliced counts obtained from the three different tools. To reduce the impact of other factors, we used the same PCA embedding and neighborhood graph for each method. For visualization, the velocities were projected onto the UMAP embedding obtained from the original processed dataset of the respective reference publication.

For the pancreatic endocrinogenesis dataset, the velocity embedding is very similar between all three methods (Fig. 4A). In particular, there is always a cyclical component in the ductal cells on the left side of the embedding and a unidirectional flow through the endocrine progenitor (EP) and pre-endocrine cells at the bottom. The only major difference lies in the β-cells, where the grid arrows in the velocyto embedding all point towards the top left corner of the cluster, while the arrows in the top left part of the cluster point towards the lower right for alevin-fry and tidesurf. On the visualization of the developing retina dataset, there are no discernible differences (Fig. S2). In contrast, the embedded velocities based on velocyto differ more strongly from those based on alevin-fry and tidesurf on the UMAPs of the 5’ datasets. For example, they are different in the DN4 and Naive clusters of the B cell data (Fig. S2B), and for the clusters c02 and c05 on the T cell data (Fig. S2C). The differences between alevin-fry and tidesurf are less pronounced.

**Fig. 4.**
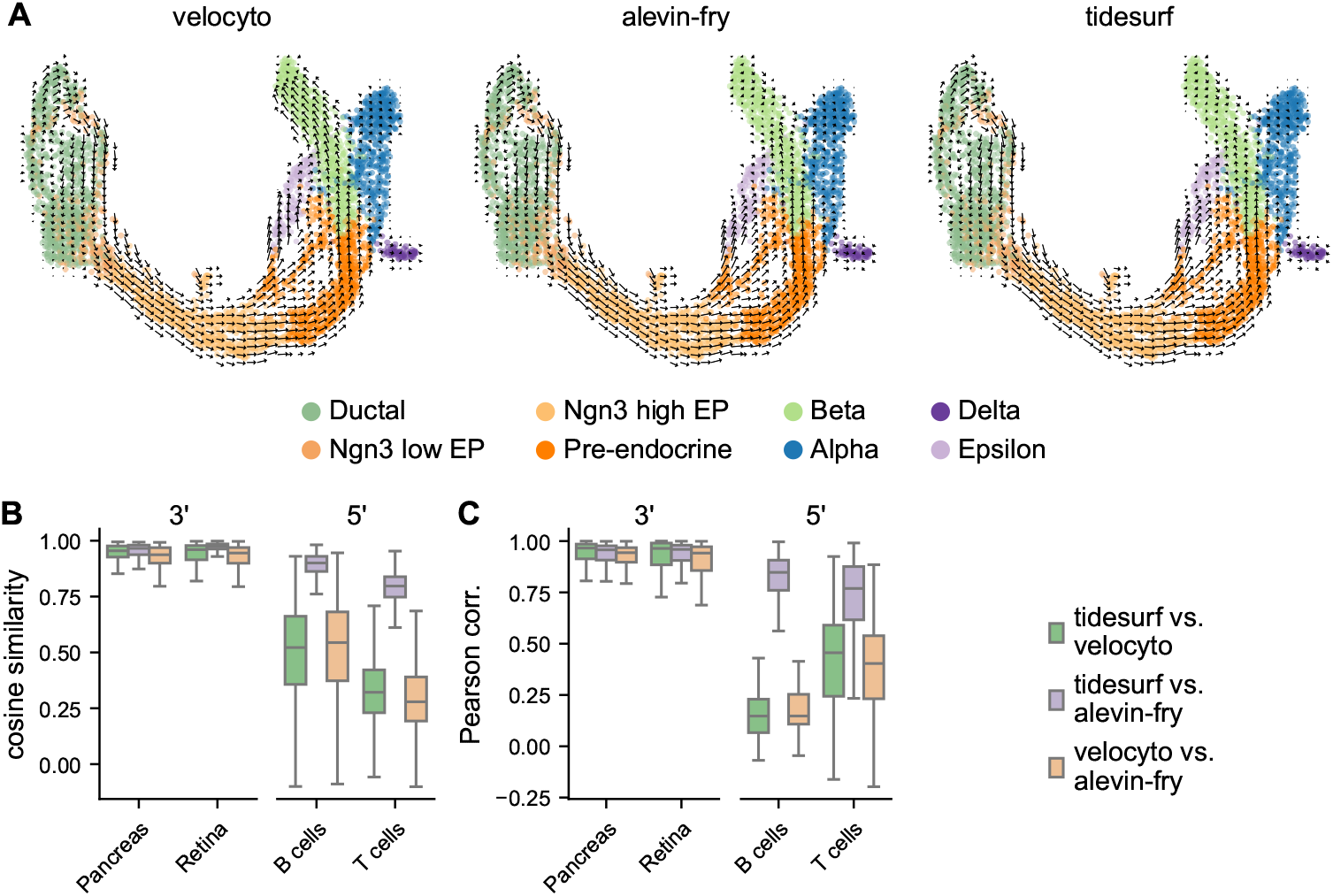
Comparison of RNA velocities between methods. **A** Visualization of velocities for the pancreatic endocrinogenesis dataset, computed from spliced and unspliced counts obtained with velocyto (left), alevin-fry (middle), or tidesurf (right) on a two-dimensional UMAP embedding. **B** Cosine similarity of velocity vectors per cell between methods for velocity genes. **C** Pearson correlation of velocities per gene between methods for velocity genes.

Additionally, we directly compared cell-wise velocity vectors. For this analysis, we focused on so-called “velocity genes”, that is, genes with a sufficiently good fit of scVelo’s model, which are used for downstream analyses including the computation of a transition probability matrix and the two-dimensional representation of velocities. Since the sets of velocity genes differed between quantification methods, we used the pairwise union of velocity genes based on the respective methods, excluding genes filtered out for either quantification based on low counts. The number of genes used in the respective pairwise comparisons as well as the number of velocity genes based on each quantification are shown in Table 2. All pairwise cosine similarities were above 0.93 in the median on the pancreas and retina datasets (Fig. 4B). For the 5’ datasets, cosine similarities were lower. Here, the highest similarities were observed between tidesurf and alevin-fry (median approximately 0.9 and 0.8 for B cells and T cells, respectively), whereas the velocities obtained based on the velocyto quantification were less similar. The median gene-wise Pearson correlation out of the pairwise union of velocity genes was above 0.93 for all comparisons on the 3’ datasets, and above 0.76 between tidesurf and alevin-fry on the 5’ data (Fig. 4C). Velocities based on velocyto had considerably lower correlations to those from the other tools (median approximately 0.15 on B cells and between 0.4 and 0.46 on T cells).

**Table 2.**
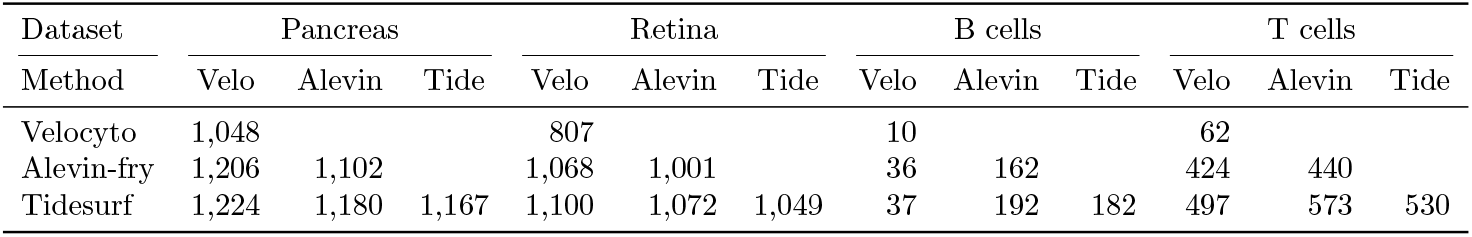
Number of velocity genes in the analyzed datasets. For each dataset, the diagonal shows the number of velocity genes based on the respective quantification. The below-diagonal elements indicate the number of genes in the pairwise union of velocity genes between the respective tools, excluding those filtered out for one of the compared quantifications because of low spliced and unspliced counts.

When comparing velocity vectors for all pairwise common genes, observed cosine similarities were considerably lower on the 5’ datasets (Fig. S3A). Even then, the similarity between alevin-fry and tidesurf was markedly higher than between velocyto and alevin-fry or tidesurf. Gene-wise Pearson correlations showed similar distributions for all genes as for velocity genes (Fig. S3B). We assume that this drop in cosine similarity is related to the well-known phenomenon that cosine similarities become more sensitive to small differences with increasing dimensionality [Blum et al., 2020]. Further, we presume that the higher differences between velocities based on alevin-fry and tidesurf, respectively, on the 5’ datasets can be attributed to the differences in unspliced counts described above as they significantly impact velocity estimation. Additionally, the B cell and T cell datasets might be less suitable for RNA velocity analysis than the pancreas and retina datasets. While the latter have been used in many RNA velocity analyses, showing biologically meaningful results, we selected the former because they had been analyzed with velocyto by the original authors, yielding erroneous velocity estimates. A strong indication of the unsuitability of these datasets for RNA velocity is their generally lower number of velocity genes (Table 2), showing that fewer genes had a good fit for scVelo’s velocity model.

### 3.4 Tidesurf has comparable run time and memory requirements as velocyto

Lastly, we also compared the run times and memory requirements of tidesurf and velocyto on a cluster with AMD EPYC 7343 CPUs. We allocated 32 cores for each process. For velocyto, the threads and memory per thread for sorting with samtools were set to 32 threads and 4 GB, respectively. To obtain accurate estimates, we selected five of the 19 samples analyzed in this study, representing the quartiles of BAM file size based on the number of reads. We executed tidesurf and velocyto five times per sample. Tidesurf had similar run times and memory requirements as velocyto (Fig. S4).

## 4 Discussion

With this work, we present tidesurf, a command line tool to quantify spliced and unspliced transcripts from 10x Genomics Chromium scRNA-seq experiments. We showed its application to four different datasets covering both 3’ and 5’ sequencing technology and demonstrated its accuracy. The accurate quantification of spliced and unspliced counts is a prerequisite for sensible RNA velocity estimates. Therefore, tidesurf represents a replacement for velocyto on 5’ data, where the latter produces highly erroneous count matrices [Klingler et al., 2025]. Additionally, we propose to use tidesurf on 3’ data as well since it generates comparable count matrices and avoids, unlike velocyto, generating an additional BAM file sorted by cells. Tidesurf’s total counts also showed considerably lower discrepancies from the Cell Ranger expression matrix than those obtained with velocyto.

In contrast to alevin-fry, tidesurf can be more easily incorporated into standard pipelines where 10x Genomics Cell Ranger is commonly used for gene expression quantification. Tidesurf directly uses Cell Ranger’s output and produces spliced, unspliced, and ambiguous counts for the same cell barcodes (or a superset if filtering is turned off) in convenient AnnData format for easy integration with Cell Ranger’s expression matrix. For alevin-fry, on the other hand, it is necessary to run a second analysis from scratch, which can lead to the calling of slightly different cell barcodes. Furthermore, the use of alevin-fry on Chromium 5’ data requires the adjustment of two parameters relating to read orientation with respect to each other and the reference genome on two different functions, which we only detected through a thorough search of multiple issues on the respective project repositories. For tidesurf, only a single command line argument has to be changed to switch from the default sense orientation of reads to reference annotations for 3’ data to the antisense orientation for 5’ data.

We compared velocity embeddings to assess the quality of tidesurf’s read count annotations as a basis for RNA velocity estimates. This comparison relied on RNA velocity estimates using scVelo’s stochastic model. We acknowledge that the resulting velocities are estimates, and that for these and their visualization shortcomings have been demonstrated [Gorin et al., 2022, Zheng et al., 2023]. The comparisons using cosine similarity and Pearson correlation offered a more detailed view and showed differences in counts and estimated velocities between the methods. However, the significance of these differences is hard to judge. Furthermore, as mentioned above, alevin-fry does not offer a simple solution for 5’ data. Therefore, we cannot exclude errors in alevin-fry’s quantifications. In the absence of a ground truth, we cannot conclusively answer the question of which quantification is more correct.

Finally, tidesurf is currently limited to 10x Genomics Chromium single-cell sequencing libraries processed with Cell Ranger. While Chromium is one of, if not the, most widely used platform for suspension-based scRNA-seq, we plan to adapt tidesurf to other single-cell protocols and alignment pipelines in future work. These adaptations would make it more widely applicable to replace velocyto.

## 5 Data availability

All analyzed datasets are publicly available and were obtained as described in subsection 2.2.

## 6 Competing interests

No competing interest is declared.

## 7 Author contributions statement

Conceptualization: J.T.S. and M.C., methodology: J.T.S., implementation: J.T.S., analysis: J.T.S., writing: J.T.S. and M.C.

## 8 Acknowledgments

We thank the members of the Claassen group for constructive discussions and feedback. Special thanks go to Matthias Bruhns and Marcello Zago for helpful feedback on code efficiency and implementation details. The authors thank the International Max Planck Research School for Intelligent Systems (IMPRS-IS) for supporting J.T.S. This work was supported by DFG CL 792/1-1.

## Supplementary Material

**Fig. S1.**
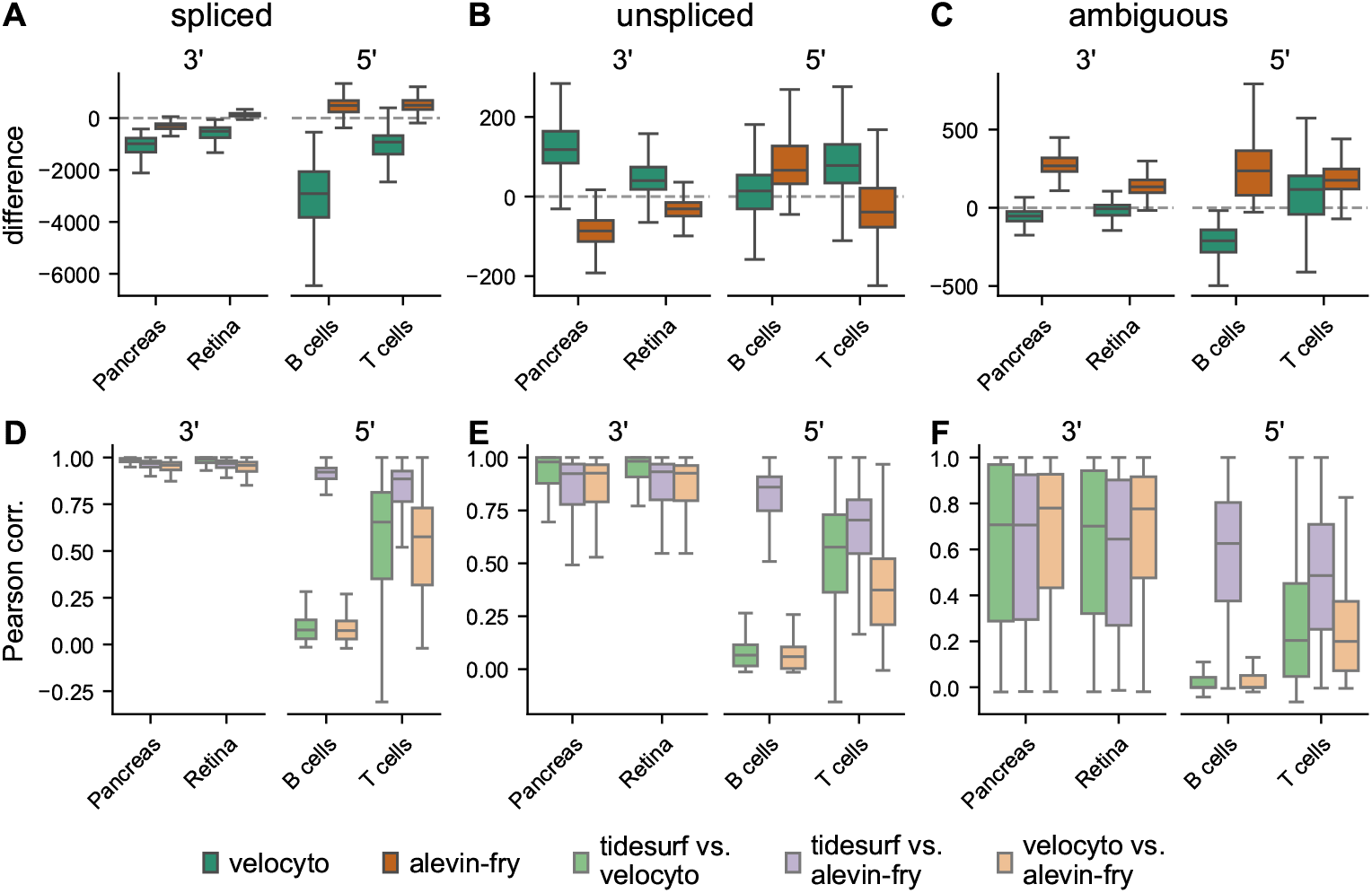
Comparison of spliced, unspliced, and ambiguous counts across methods. **A-C** Difference of total spliced (**A**), unspliced (**B**), and ambiguous (**C**) counts per cell from tidesurf quantification. **D-F** Pearson correlation of spliced (**D**), unspliced (**E**), and ambiguous (**F**) counts per gene between methods.

**Fig. S2.**
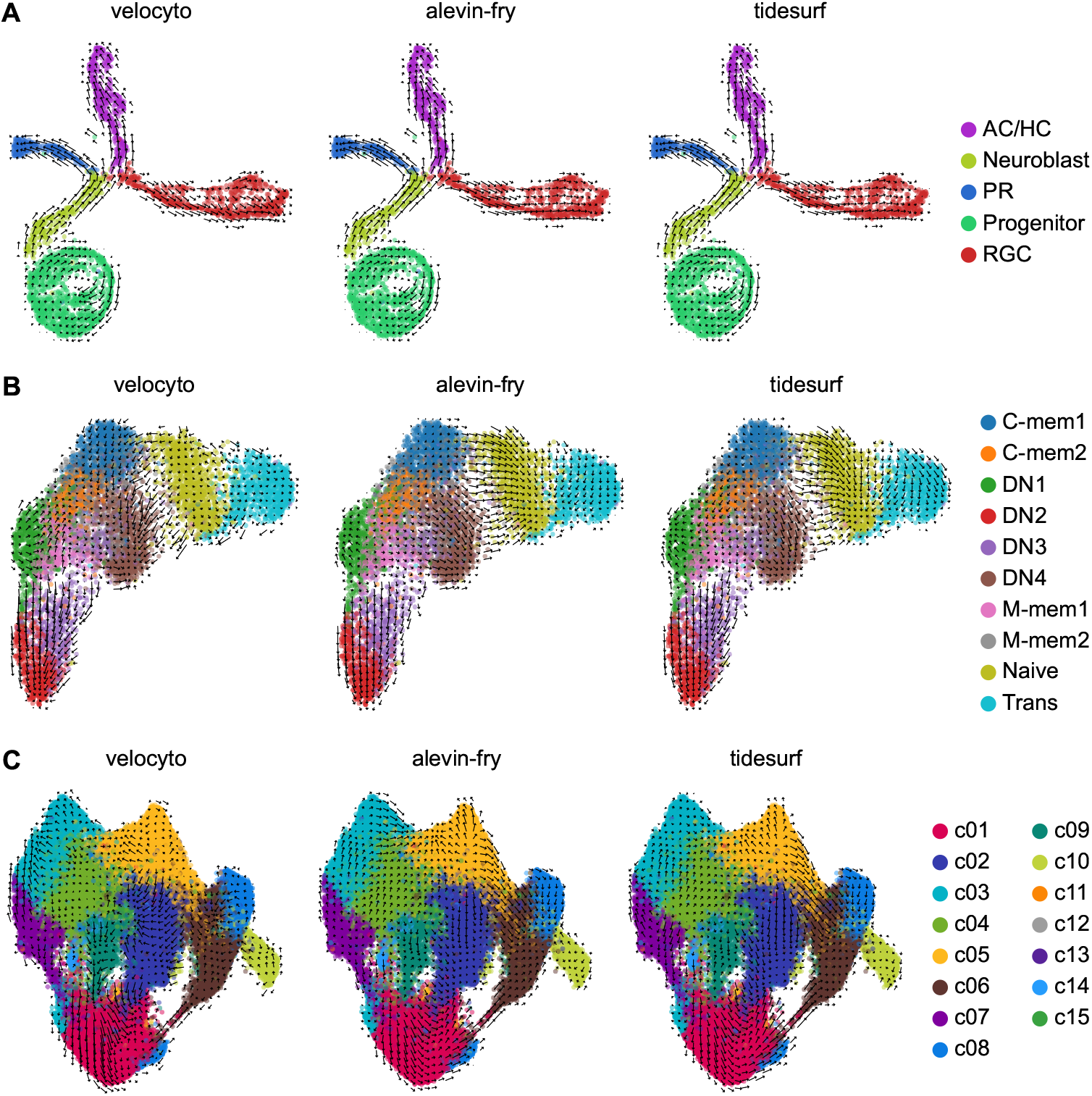
Visualization of RNA velocities based on different tools. **A-C** Visualization of velocities for the retina (**A**), B cell (**B**), and T cell (**C**) datasets, computed from spliced and unspliced counts obtained with velocyto (left panels), alevin-fry (middle panels), or tidesurf (right panels) on a two-dimensional UMAP embedding of the respective dataset.

**Fig. S3.**
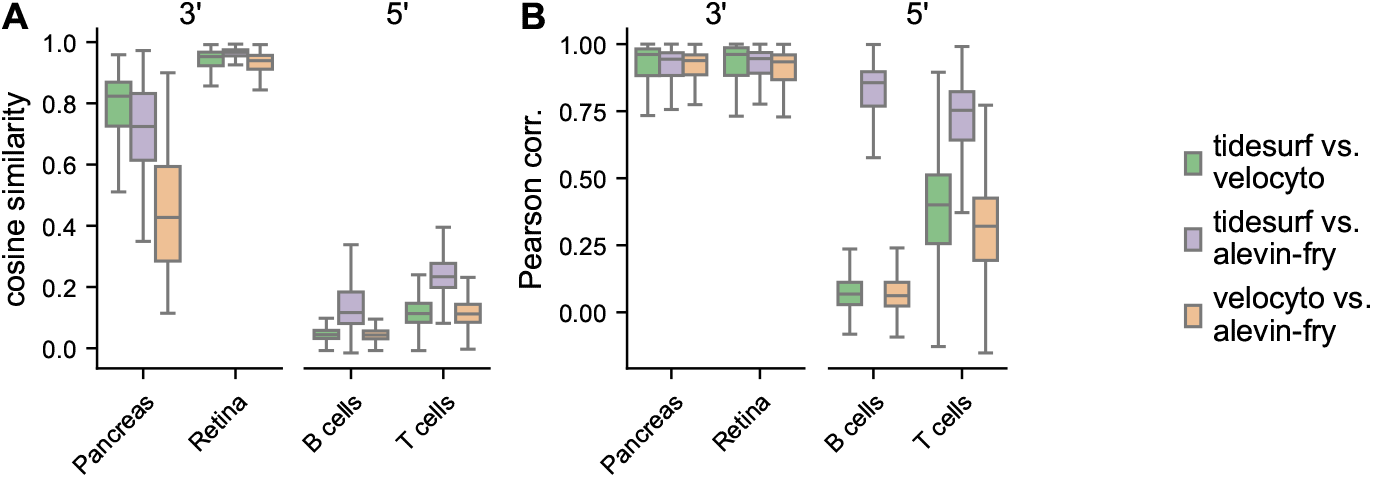
Comparison of RNA velocities between methods. **A** Cosine similarity of velocity vectors per cell between methods for all common genes. **B** Pearson correlation of velocities per gene between methods for all common genes.

**Fig. S4.**
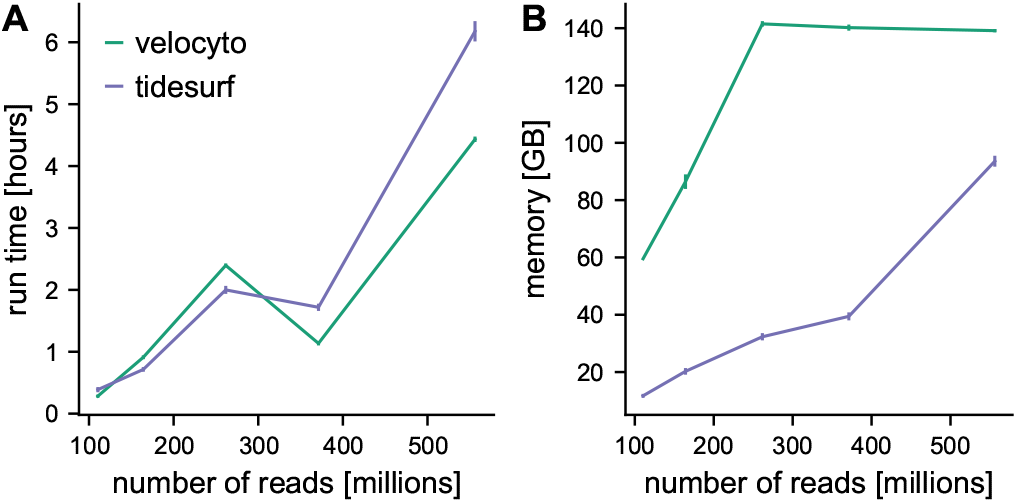
Comparison of run time and memory requirements. **A-B** Mean run time (**A**) and utilized memory (**B**) for five selected samples, plotted against the number of reads in the sample. The lines represent the mean of five runs. Error bars show standard deviation.

## Notes

### Competing Interest Statement

The authors have declared no competing interest.

## References

0x Genomics. Cell Ranger’s Gene Expression Algorithm. https://www.10xgenomics.com/support/software/cell-ranger/latest/algorithms-overview/cr-gex-algorithm, 2024. [Online; accessed 27.12.2024].

A. Bastidas-Ponce, S. Tritschler, L. Dony, K. Scheibner, M. Tarquis-Medina, C. Salinno, S. Schirge, I. Burtscher, A. Böttcher, F. J. Theis, H. Lickert, M. Bakhti, A. Klein, and B. Treutlein. Comprehensive single cell mRNA profiling reveals a detailed roadmap for pancreatic endocrinogenesis. Development, 146(12):dev173849, 06 2019. ISSN 0950-1991. doi: 10.1242/dev.173849. URL https://doi.org/10.1242/dev.173849.

V. Bergen, M. Lange, S. Peidli, F. A. Wolf, and F. J. Theis. Generalizing RNA velocity to transient cell states through dynamical modeling. Nature Biotechnology, 38(12): 1408–1414, 2020. ISSN 1546-1696. doi: 10.1038/s41587-020-0591-3. URL https://doi.org/10.1038/s41587-020-0591-3.

A. Blum, J. Hopcroft, and R. Kannan. Foundations of Data Science. Cambridge University Press, Cambridge, 2020. ISBN 9781108485067. doi: 10.1017/9781108755528. URL https://doi.org/10.1017/9781108755528.

A. Dobin, C. A. Davis, F. Schlesinger, J. Drenkow, C. Zaleski, S. Jha, P. Batut, M. Chaisson, and T. R. Gingeras. STAR: ultrafast universal RNA-seq aligner. Bioinformatics, 29(1):15–21, 10 2012. ISSN 1367-4803. doi: 10.1093/bioinformatics/bts635. URL https://doi.org/10.1093/bioinformatics/bts635.

J. Fu, Z. Wang, M. Martinez, A. Obradovic, W. Jiao, K. Frangaj, R. Jones, X. V. Guo, Y. Zhang, W.-I. Kuo, H. M. Ko, A. Iuga, C. Bay Muntnich, A. Prada Rey, K. Rogers, J. Zuber, W. Ma, M. Miron, D. L. Farber, J. Weiner, T. Kato, Y. Shen, and M. Sykes. Plasticity of intragraft alloreactive T cell clones in human gut correlates with transplant outcomes. Journal of Experimental Medicine, 221(1):e20230930, 12 2023. ISSN 0022-1007. doi: 10.1084/jem.20230930. URL https://doi.org/10.1084/jem.20230930.

M. Gao, C. Qiao, and Y. Huang. UniTVelo: temporally unified RNA velocity reinforces single-cell trajectory inference. Nature Communications, 13(1):6586, Nov 2022. ISSN 2041-1723. doi: 10.1038/s41467-022-34188-7. URL https://doi.org/10.1038/s41467-022-34188-7.

G. Gorin, M. Fang, T. Chari, and L. Pachter. RNA velocity unraveled. PLOS Computational Biology, 18(9):1–55, 09 2022. doi: 10.1371/journal.pcbi.1010492. URL https://doi.org/10.1371/journal.pcbi.1010492.

J. A. Griffiths, A. Scialdone, and J. C. Marioni. Using single-cell genomics to understand developmental processes and cell fate decisions. Molecular Systems Biology, 14(4):e8046, 2018. doi: 10.15252/msb.20178046. URL https://www.embopress.org/doi/abs/10.15252/msb.20178046.

D. He, M. Zakeri, H. Sarkar, C. Soneson, A. Srivastava, and R. Patro. Alevin-fry unlocks rapid, accurate and memory-frugal quantification of single-cell RNA-seq data. Nature Methods, 19(3):316–322, 2022. ISSN 1548-7105. doi: 10.1038/s41592-022-01408-3. URL https://doi.org/10.1038/s41592-022-01408-3.

D. Klingler, J. T. Schleicher, and M. Claassen. Quantification of spliced and unspliced transcripts by velocyto is inaccurate for 5’-sequencing data. bioRxiv, 2025. doi: 10.1101/2025.01.17.633503. URL https://www.biorxiv.org/content/early/2025/01/22/2025.01.17.633503.

G. La Manno, R. Soldatov, A. Zeisel, E. Braun, H. Hochgerner, V. Petukhov, K. Lidschreiber, M. E. Kastriti, P. Lönnerberg, A. Furlan, J. Fan, L. E. Borm, Z. Liu, D. van Bruggen, J. Guo, X. He, R. Barker, E. Sundström, G. Castelo-Branco, P. Cramer, I. Adameyko, S. Linnarsson, and P. V. Kharchenko. RNA velocity of single cells. Nature, 560(7719):494–498, 2018. ISSN 1476-4687. doi: 10.1038/s41586-018-0414-6. URL https://doi.org/10.1038/s41586-018-0414-6.

Q. Lo Giudice, M. Leleu, G. La Manno, and P. J. Fabre. Single-cell transcriptional logic of cell-fate specification and axon guidance in early-born retinal neurons. Development, 146(17):dev178103, 09 2019. ISSN 0950-1991. doi: 10.1242/dev.178103. URL https://doi.org/10.1242/dev.178103.

J. Y. Lu, W. Shao, L. Chang, Y. Yin, T. Li, H. Zhang, Y. Hong, M. Percharde, L. Guo, Z. Wu, L. Liu, W. Liu, P. Yan, M. Ramalho-Santos, Y. Sun, and X. Shen. Genomic repeats categorize genes with distinct functions for orchestrated regulation. Cell Reports, 30(10):3296–3311.e5, Mar 2020. ISSN 2211-1247. doi: 10.1016/j.celrep.2020.02.048. URL https://doi.org/10.1016/j.celrep.2020.02.048.

P. Melsted, A. S. Booeshaghi, L. Liu, F. Gao, L. Lu, K. H. J. Min, E. da Veiga Beltrame, K. E. Hjörleifsson, J. Gehring, and L. Pachter. Modular, efficient and constant-memory single-cell RNA-seq preprocessing. Nature Biotechnology, 39(7): 813–818, Jul 2021. ISSN 1546-1696. doi: 10.1038/s41587-021-00870-2. URL https://doi.org/10.1038/s41587-021-00870-2.

R. Nayak and Y. Hasija. A hitchhiker’s guide to single-cell transcriptomics and data analysis pipelines. Genomics, 113(2):606–619, 2021. ISSN 0888-7543. doi: 10.1016/j.ygeno.2021.01.007. URL https://www.sciencedirect.com/science/article/pii/S0888754321000331.

R. Patro, G. Duggal, M. I. Love, R. A. Irizarry, and C. Kingsford. Salmon provides fast and bias-aware quantification of transcript expression. Nature Methods, 14(4): 417–419, Apr 2017. ISSN 1548-7105. doi: 10.1038/nmeth.4197. URL https://doi.org/10.1038/nmeth.4197.

A. Stewart, J. C.-F. Ng, G. Wallis, V. Tsioligka, F. Fraternali, and D. K. Dunn-Walters. Single-Cell Transcriptomic Analyses Define Distinct Peripheral B Cell Subsets and Discrete Development Pathways. Frontiers in Immunology, 12, 2021. ISSN 1664-3224. doi: 10.3389/fimmu.2021.602539. URL https://www.frontiersin.org/journals/immunology/articles/10.3389/fimmu.2021.602539.

I. Virshup, S. Rybakov, F. J. Theis, P. Angerer, and F. A. Wolf. anndata: Access and store annotated data matrices. Journal of Open Source Software, 9(101):4371, 2024. doi: 10.21105/joss.04371. URL https://doi.org/10.21105/joss.04371.

F. A. Wolf, P. Angerer, and F. J. Theis. SCANPY: large-scale single-cell gene expression data analysis. Genome Biology, 19(1):15, Feb 2018. ISSN 1474-760X. doi: 10.1186/s13059-017-1382-0. URL https://doi.org/10.1186/s13059-017-1382-0.

G. X. Y. Zheng, J. M. Terry, P. Belgrader, P. Ryvkin, Z. W. Bent, R. Wilson, S. B. Ziraldo, T. D. Wheeler, G. P. McDermott, J. Zhu, M. T. Gregory, J. Shuga, L. Montesclaros, J. G. Underwood, D. A. Masquelier, S. Y. Nishimura, M. Schnall-Levin, P. W. Wyatt, C. M. Hindson, R. Bharadwaj, A. Wong, K. D. Ness, L. W. Beppu, H. J. Deeg, C. McFarland, K. R. Loeb, W. J. Valente, N. G. Ericson, E. A. Stevens, J. P. Radich, T. S. Mikkelsen, B. J. Hindson, and J. H. Bielas. Massively parallel digital transcriptional profiling of single cells. Nature Communications, 8(1):14049, Jan 2017. ISSN 2041-1723. doi: 10.1038/ncomms14049. URL https://doi.org/10.1038/ncomms14049.

S. C. Zheng, G. Stein-O’Brien, L. Boukas, L. A. Goff, and K. D. Hansen. Pumping the brakes on RNA velocity by understanding and interpreting RNA velocity estimates. Genome Biology, 24(1):246, Oct 2023. ISSN 1474-760X. doi: 10.1186/s13059-023-03065-x. URL https://doi.org/10.1186/s13059-023-03065-x.

